# *C9ORF72* dipeptide repeat proteins disrupt formation of GEM bodies and induce aberrant accumulation of survival of motor neuron protein

**DOI:** 10.1101/2021.03.24.436890

**Authors:** Yuma Kato, Minnie Yokogawa, Ikuma Nakagawa, Kazunari Onodera, Hideyuki Okano, Haruhisa Inoue, Mitsuharu Hattori, Yohei Okada, Hitomi Tsuiji

## Abstract

A GGGGCC repeat expansion in the *C9ORF72* gene is the most common genetic cause of amyotrophic lateral sclerosis (ALS), a devastating motor neuron disease. In the neurons of ALS patients, dipeptide repeat proteins (DPRs) are produced from repeat-containing RNAs by an unconventional form of translation, and some of these proteins, especially those containing poly(glycine-arginine) and poly(proline-arginine), are toxic to neurons. Gemini of coiled bodies (GEMs) are nuclear structures that harbor survival of motor neuron (SMN) protein, and SMN is essential for the assembly of U-rich small nuclear ribonucleoproteins (snRNPs) that are central for splicing. We previously reported that GEMs are lost and that snRNP biogenesis is misregulated in the motor neurons of ALS patients. Here we show that DPRs interfere with GEM formation and proper SMN localization in HeLa cells and iPSC-derived motor neurons from an ALS patient with the *C9ORF72* mutation. The accumulation of poly(glycine-arginine) markedly reduced the number of GEMs and caused the formation of aberrant cytoplasmic RNA granules that sequestered SMN. These findings indicate the functional impairment of SMN in motor neurons expressing DPRs and may provide a mechanism to explain the vulnerability of motor neurons of C9ORF72-ALS patients.

## Introduction

Amyotrophic lateral sclerosis (ALS) is a late-onset devastating motor neuron disease, characterized by progressive loss of motor neurons (MNs) resulting in muscle weakness and atrophy ^1^. A large expansion of a GGGGCC nucleotide repeat in the first intron of the *C9ORF72* gene is the most common genetic cause of ALS and frontotemporal dementia (FTD) (C9-ALS/FTD) ^2,3^. Both sense and antisense strands of the expanded repeat are transcribed, and the resulting RNAs are translated by an unconventional form of translation called repeat-associated non-AUG (RAN) translation to produce dipeptide repeat (DPR) proteins ^4–6^. These DPR proteins accumulate in the affected neurons of patients with *C9ORF72* mutations and represent a pathological hallmark. DPRs are cytotoxic in cells and multiple animal models ^7–10^. Among them, the arginine-rich ones poly(glycine-arginine, GR) and poly(proline-arginine, PR) are the most toxic to neurons ^7–9^. The pattern of poly(GR) accumulation seems to be the most correlated with neurodegeneration ^11,12^.

On the other hand, spinal muscular atrophy (SMA) is early-onset motor neuron disease, caused by the loss of survival of motor neuron (SMN) protein, due to mutation in the *SMN1* gene ^13–16^. SMN is essential for the biogenesis of U-rich small nuclear ribonucleoproteins (snRNPs), which are the major components of the spliceosome, the machinery that carries out pre-mRNA splicing ^17^. Accordingly, there is a tight correlation between the degree of SMN deficiency and snRNP assembly in SMA ^17,18^. In vertebrates, GEMs (Gemini of coiled bodies) consist of SMN, seven Gemin proteins (Gemin 2–Gemin 8), and Unrip. The precise function of GEMs is still unclear, although the amount of SMN protein correlates well with the number of GEMs, and the number of GEMs is considered to be a good indicator of SMA severity ^19^. snRNPs consist of one or two small nuclear RNAs (snRNAs) bound to a set of seven Smith (Sm) or Smith-like proteins and a unique set of snRNP-specific proteins ^20^. U1, U2, U4/U6, and U5 snRNPs form the major spliceosome, which is responsible for splicing of a majority of pre-mRNA introns ^21^. In SMA patients, SMN deficiency causes abnormal snRNP biogenesis and widespread defects in splicing that might be the major causes of neurodegeneration ^18^.

We and others have previously found that GEMs, the site of SMN protein localization in the nucleus, are lost and that snRNP biogenesis is dysregulated in affected MNs of patients with ALS and SMA ^22,23^. Loss of GEMs and/or abnormal snRNP biogenesis has also been observed in fibroblasts of ALS patients harboring TDP-43 or FUS mutations ^24^ and in various ALS mouse models expressing mutant SOD1 or TDP-43 ^25,26^. Enhancing the level of SMN protein attenuates neurodegeneration in a mouse model of SMA, transgenic mutant TDP-43 mice ^27^ and mutant SOD1 mice ^28^. SMN directly binds to FUS, another RNA-binding protein associated with ALS, providing further evidence for the link between ALS and SMA ^22,24,29^. These findings underscore the importance of SMN protein and proper snRNP biogenesis in ALS pathogenesis. However, the direct causes of loss of GEMs and abnormal snRNP biogenesis in ALS patients have been obscure.

In this study, we show that poly(GR) and poly(PR) affect the localization of SMN and result in the loss of GEMs and the cytoplasmic aggregation of SMN and RNA-binding proteins. We also provide evidence for decreased number of GEMs in MNs differentiated from iPSCs of an ALS patient with a *C9ORF72* mutation.

## Results

### Poly(GR) disrupts GEM formation

We first investigated the effect of *C9ORF72* poly(GR) and poly(PR) on GEM formation. We transfected HeLa cells with a plasmid encoding a FLAG-tagged construct with 50 repeats of glycine-arginine (poly(GR)) or a FLAG-tagged construct with 50 repeats of proline-arginine (poly(PR)). We detected GEMs by immunostaining for SMN protein and detected DPRs using an anti-FLAG antibody. Both poly(GR) and poly(PR) localized to the nucleolus (Fig. 1a), consistent with previous reports ^7^, whereas poly(GR) also distributed in the cytoplasm very weakly (data not shown). In HeLa cells transfected with an empty vector, we observed approximately 8 GEMs per cell (Fig. 1a, upper row,1B). In cells with a poly(GR) accumulation in the nucleolus, the number of GEMs was reduced to approximately 2 (Fig. 1a, middle row,1b). Poly(PR) accumulation in the nucleolus considerably reduced the number of GEMs, but to a lesser extent than poly(GR) and not in a statistically significant way (Fig. 1a, bottom row, 1b, p = 0.0768). These observations clearly indicate that DPRs especially poly(GR) disrupts the formation of GEMs in nucleus.

**Figure 1.**
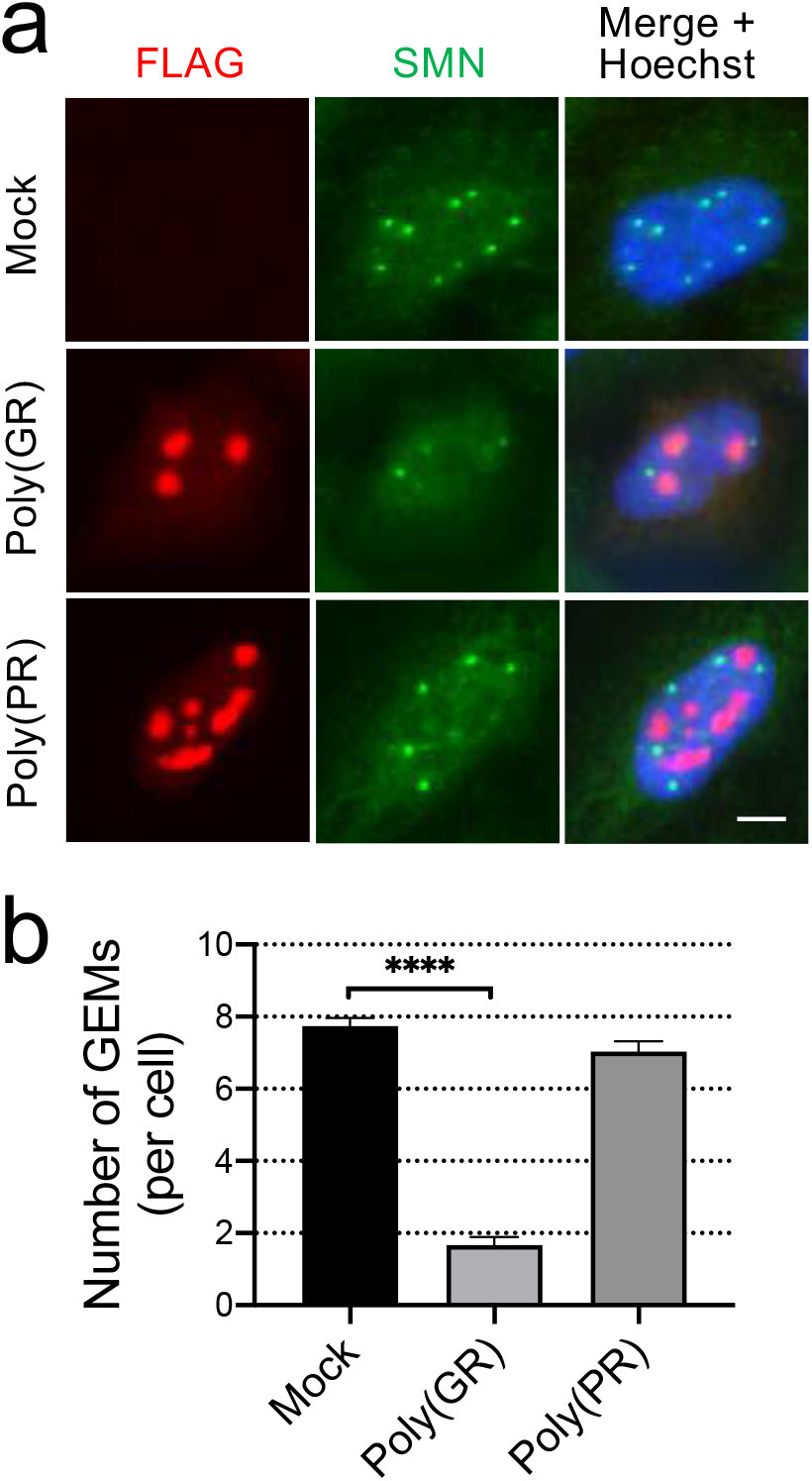
C9ORF72 poly(GR) disrupts GEM formation. (a) HeLa-C3 cells were transfected with empty vector (mock), a FLAG-tagged construct expressing 50 repeats of GR (poly(GR)) or a FLAG-tagged construct expressing 50 repeats of PR (poly(PR)) and immunostained with anti-FLAG and anti-SMN antibodies. SMN-positive nuclear granules are GEMs. Nuclei were counterstained with Hoechst. Scale bar = 10 μm. (b) (b)Quantification of the number of GEMs visualized in (A). Cells transfected with empty vector (mock), cells with an accumulation of poly(GR) in the nucleolus, and cells with poly(PR) were analysed. More than 100 cells were counted. Biological replicates; n = 3~5. One-way ANOVA with Tukey’s multiple comparisons tests, ****; p <0.0001.

### Poly(GR) and poly(PR) induce cytoplasmic RNA granules harboring SMN protein

We also noticed that SMN and poly(GR) occasionally accumulated in the cytoplasm of cells expressing poly(GR). Therefore, we next investigated whether DPRs induce abnormal cytoplasmic granules. Poly(A)-binding protein (PABP) was distributed diffusely in the cytoplasm of cells with mock transfection, whereas PABP formed foci resembling stress granules in the cytoplasm of cells expressing poly(GR) (Fig. 2a). We observed cytoplasmic accumulation of PABP in approximately 80% of cells with poly(GR) expression in the nucleolus (Fig. 2b). Some PABP accumulations harbored poly(GR) (Fig. 2a), and SMN occasionally colocalized with poly(GR) or PABP in the cytoplasm (Fig. 2c, 2d). These PABP- and poly(GR)-containing cytoplasmic granules may also contain RNA, which we visualized by SYTO RNASelect Green Fluorescent Cell Stain (Fig. 2e). Poly(PR) also led to the formation of PABP granules resembling stress granules but to a lesser extent than poly(GR) (Fig. 2a, bottom row, 2b, p = 0.0158). Notably, poly(PR) did not colocalize with PABP in cytoplasmic granules induced by poly(PR) (Fig. 2a, bottom row), though SMN occasionally localized in the cytoplasmic PABP granules (Fig. 2b, 2c). Intriguingly, the average number of GEMs was reduced to 1.23 in the cells with SMN accumulation in the cytoplasm (n = 3, more than 10 cells were counted). Together, these findings suggest that *C9ORF72* DPRs can disrupt GEMs and cause mislocalization of SMN protein to the cytoplasm, where they accumulate with RNA protein granules containing PABP.

**Figure 2.**
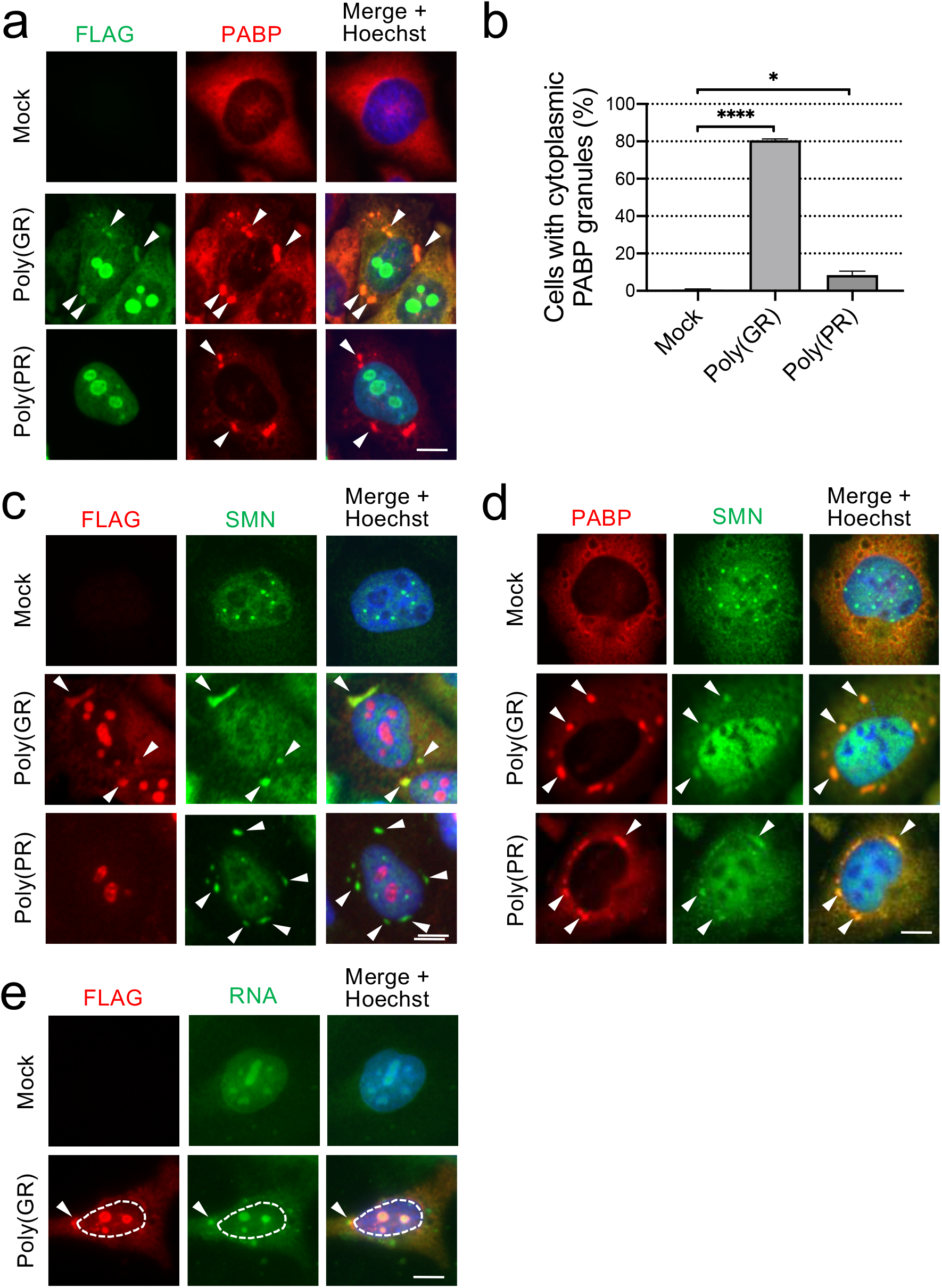
C9ORF72 poly(GR) and poly(PR) induce cytoplasmic RNA granules harboring SMN protein. (a) HeLa-C3 cells were transfected with empty vector (mock), FLAG-poly(GR), or FLAG-poly(PR) and immunostained with anti-FLAG and anti-PABP antibodies.PABP was distributed diffusely in the cytoplasm of cells with mock transfection, whereas PABP formed foci in the cytoplasm of cells with poly(GR) or poly(PR). Notably, PABP-granules contained poly(GR) but not poly(PR). Nuclei were counterstained with Hoechst. Cytoplasmic granules are indicated with arrowheads. Scale bar = 10 μm. (b) HeLa-C3 cells were transfected with empty vector (mock), FLAG-poly(GR), or FLAG-poly(PR) and immunostained with anti-FLAG and anti-SMN antibodies. SMN formed foci in the cytoplasm of cells with poly(GR) or poly(PR). SMN occasionally colocalized with poly(GR) in the cytoplasm. Nuclei were counterstained with Hoechst. Cytoplasmic granules are indicated with arrowheads. Scale bar = 10 μm. (c) HeLa-C3 cells were transfected with empty vector (mock), FLAG-poly(GR), or FLAG-poly(PR) and immunostained with anti-SMN and anti-PABP antibodies. SMN occasionally colocalized with PABP. Nuclei were counterstained with Hoechst. Cytoplasmic granules are indicated with arrowheads. Scale bar = 10 μm. (d) HeLa-C3 cells were transfected with empty vector (mock) or FLAG-poly(GR), and RNA was visualized with SYTO RNAselect (Molecular Probes). Nuclei were counterstained with Hoechst. Cytoplasmic granules are indicated with arrowheads. Scale bar = 10 μm. (e) Quantification of the number of cytoplasmic PABP granules visualized in (a). Cells transfected with empty vector (mock), cells with an accumulation of poly(GR) in the nucleolus, and cells with poly(PR) were analysed. More than 100 cells were counted. Biological replicates; n= 3~5. One-way ANOVA with Tukey’s multiple comparisons tests, *; p <0.05, ****; p <0.0001.

### Poly(PR) expression combined with oxidative stress induces persistent cytoplasmic stress granules harboring SMN

Based on the results of these experiments in which poly(GR) and poly(PR) caused the formation of cytoplasmic PABP granules resembling stress granules, and SMN was inactivated by oxidative stress 30, we next tested whether stress granule formation could induce loss of GEMs. HeLa cells were treated with 500 μM sodium arsenite for 30 min and immunostained with anti-PABP and anti-SMN antibodies. The formation of stress granules itself did not induce the loss of GEMs (Supplementary Fig. S1), indicating that sequestration of SMN in stress granules is not sufficient to induce the loss of GEMs. We further tested the combined effects of DPR toxicity and oxidative stress on SMN. We treated HeLa cells with 500 μM sodium arsenite for 30 min to induce oxidative stress and PABP-containing stress granule formation. Following arsenite treatment, we washed the cells to remove the arsenite and allowed the cells to recover and the stress granules to disassemble. In control cells, the stress granules disassembled almost completely, and PABP staining was dispersed 3 hours postincubation. However, in cells expressing poly(PR), cytoplasmic granules became more persistent and remained in the cytoplasm even after 6 hours (Fig. 3a, 3b). This is consistent with other reports showing that DPRs can induce persistent stress granule formation ^31,32^. These persistent cytoplasmic granules occasionally harbored SMN protein (Fig. 3b), and the average number of GEMs was reduced to 0.7 in these cells (n = 3, more than 10 cells were counted). This sequestration of SMN in abnormal cytoplasmic granules could disturb the normal function of SMN (illustrated in Fig. 5).

**Figure 3.**
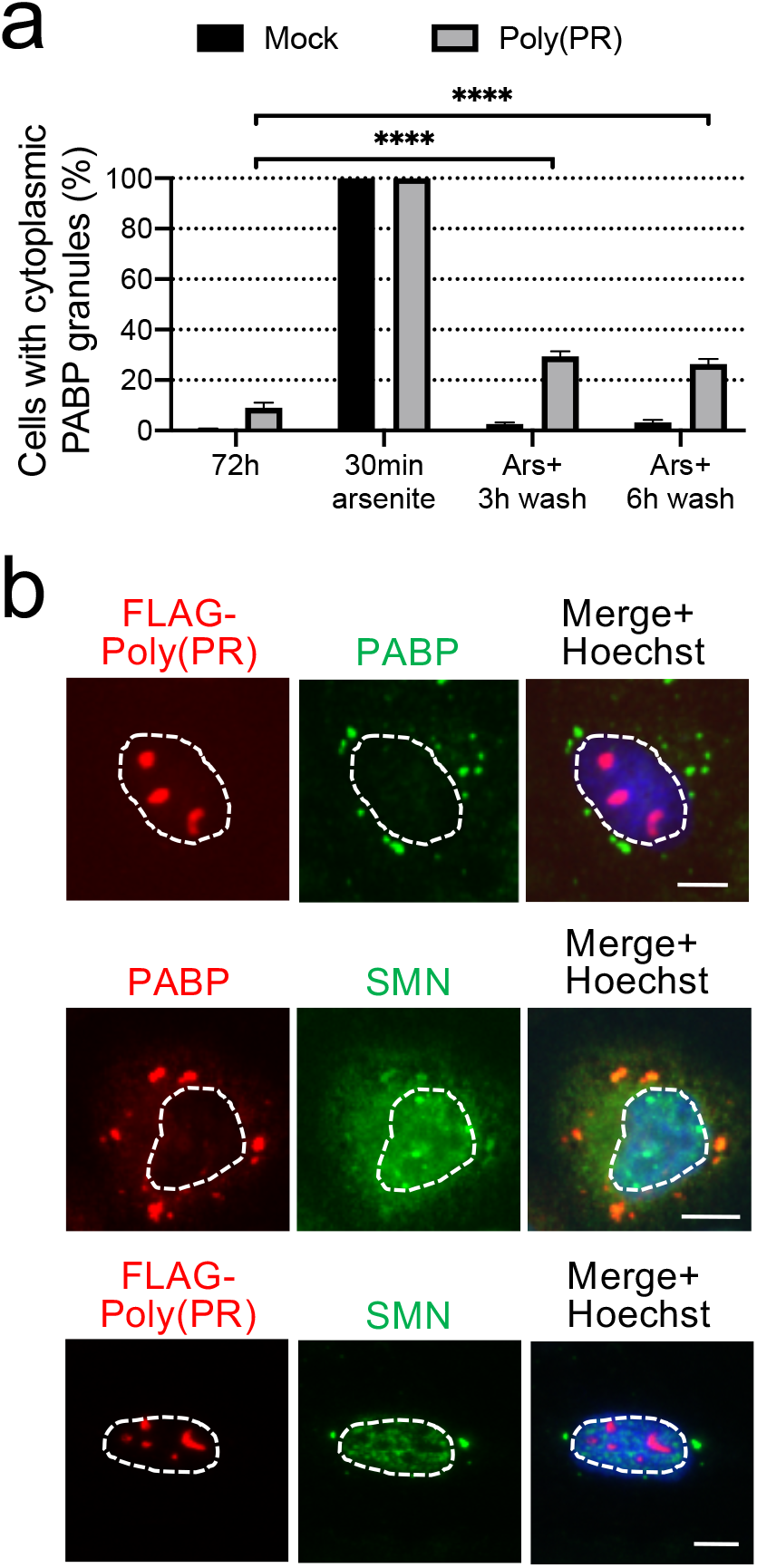
Persistent cytoplasmic stress granules induced by poly(PR) and oxidative stress sequester SMN. (a) Quantification of the number of cytoplasmic stress granules induced by poly(PR) and oxidative stress. HeLa-C3 cells were treated with 500 μM sodium arsenite for 30 min to induce cytoplasmic stress granules. Cells were then washed to remove the arsenite and allowed to recover and the stress granules to disassemble. Cells were fixed at the indicated time points and immunostained with anti-FLAG and anti-PABP antibodies. Cells with cytoplasmic PABP granules were counted. More than 100 cells were counted. Biological replicates; n = 3~5. Two-way ANOVA with Tukey’s multiple comparisons tests, ****; p <0.0001. (b) Immunostained images of persistent cytoplasmic granules induced in (a). Cells were immunostained with anti-FLAG, anti-PABP, and anti-SMN antibodies. Scale bar = 10 μm. Nuclei counterstained with Hoechst are enclosed by dotted lines.

### The number of GEMs in MNs derived from iPSCs of a C9ORF72-ALS patient is reduced

We previously showed that GEMs are almost completely lost in MNs in the spinal cord of postmortem ALS patients ^22^. To provide insight whether C9ORF72 mutation affect GEM pathology in patient-derived MNs, we took advantage of human iPSCs derived from an ALS patient with a *C9ORF72* mutation (C9-ALS) and two healthy controls. The origins of the iPSC clones used in this analysis are described in Table 1. Differentiation of iPSCs into MNs was performed as previously described ^33–35^ (Fig. 4a). The differentiation efficiency into MNs was approximately the same in all three iPSCs used in this study, as evidenced by HB9 and Isl-1 immunostaining at day 7 of adherent differentiation (Fig. 4b). MNs differentiated from iPSCs derived from an ALS patient with a *C9ORF72* mutation (C9-ALS) or control patients were fixed for the analysis at 4 weeks of adherent differentiation. iPSC-derived MNs were visualized by the lentivirus reporter specific for motor neurons (*HB9^e438^::Venus*). To count the number of GEMs, we immunostained MNs with anti-SMN and anti-Gemin-2 antibodies, and SMN-positive and Gemin-2-positive granules in the nuclei were counted as GEMs. We observed a decreased number of GEMs in MNs differentiated from iPSCs of the ALS patient compared with that in MNs from healthy donors (Fig. 4c, p = 0.0002 (Contol1 vs C9-ALS) and = 0.0002 (Contol2 vs C9-ALS), 4d).

**Table 1.** List of iPSC clones.

|  | clone name<br>at<br>establishment | gender | onset<br>age<br>(y) | age at<br>diagnosis | biopsy<br>age<br>(y) | origin | reprogramming | references |
| --- | --- | --- | --- | --- | --- | --- | --- | --- |
| Control 1 | TIGE-9 | male | N.A. | N.A. | 36 | skin fibroblasts | episomal | Shimojo et al., 2015 <sup>33</sup> |
| Control 2 | 201B7 | female | N.A. | N.A. | 36 | skin fibroblasts | retrovirus | Takahashi et al., 2007 <sup>48</sup> |
| C9-ALS | ND06769E4 | female | 45 | 46 | 46 | immortalized<br>B-lymphocytes | episomal | Imamura et al., 2017 <sup>49</sup> |

**Figure 4.**
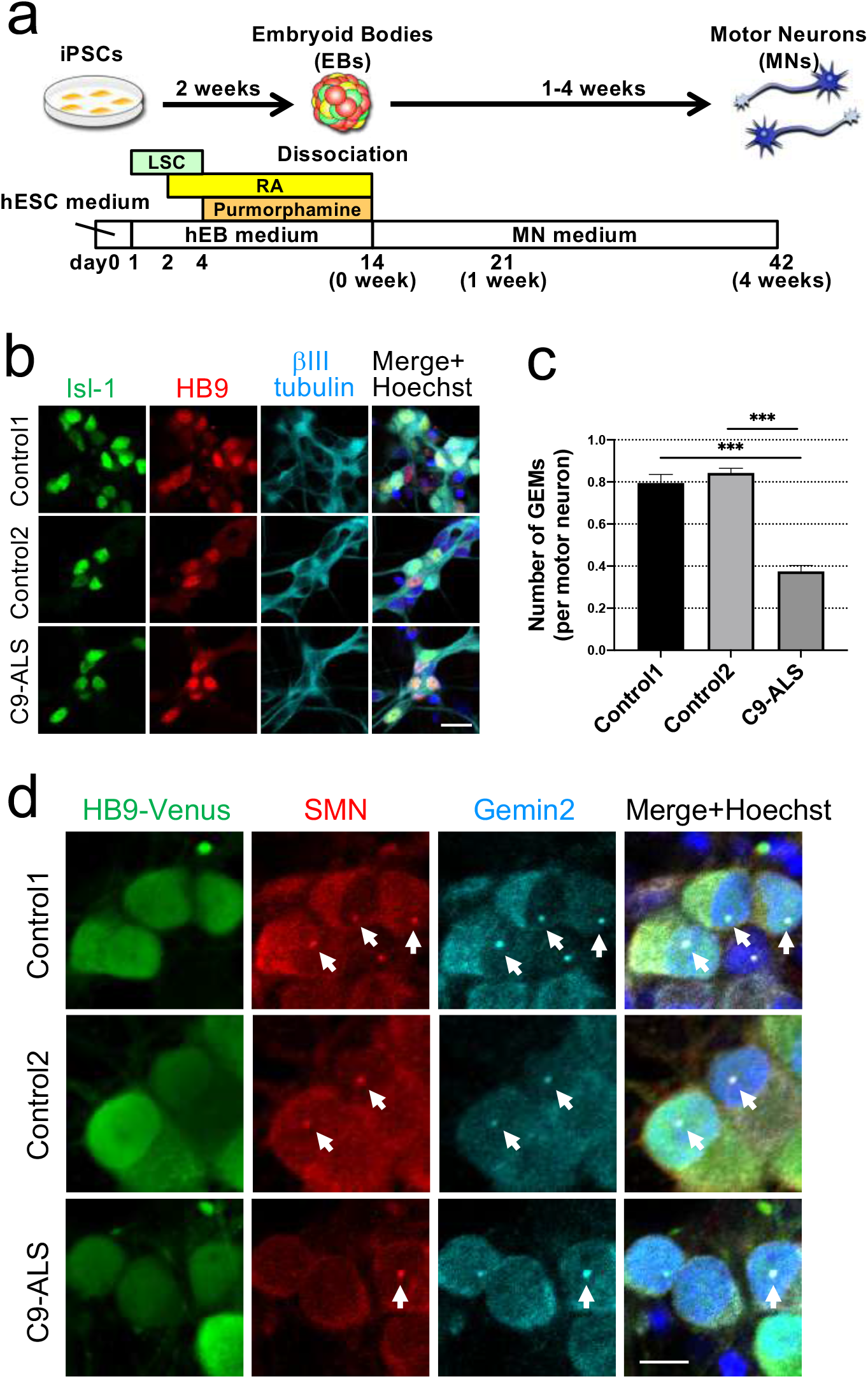
The number of GEMs in MNs derived from iPSCs of a C9ORF72-ALS patient is reduced. (a) Schematic of the protocol for the induction of MNs from iPSCs of healthy subjects and a C9ORF72-ALS patient ^33^. LSC: LDN-193189, SB431542, and CHIR99021, RA: retinoic acid. (b) Differentiation of MNs from iPSCs. Cells were fixed and analysed with anti-Isl-1, anti-HB9, and anti-βIII tubulin antibodies on day 7 of adherent differentiation. The differentiation efficiencies for the three iPSCs were approximately the same. Nuclei were counterstained with Hoechst. Scale bar = 20 μm. (c) Quantification of the number of GEMs in MNs differentiated from iPSCs of healthy donors and a C9ORF72-ALS patient. MNs were stained with anti-GFP (HB9^e436^::Venus), anti-SMN, and anti-Gemin 2 antibodies at 4 weeks of differentiation. MNs were identified by HB9^e436^::Venus expression. SMN- and Gemin 2-double positive granules were counted as GEMs. More than 100 cells were counted. Biological replicates; n = 3. One-way ANOVA with Tukey’s multiple comparisons tests, ***; p <0.001. (d) Representative confocal images of GEMs in MNs differentiated from iPSCs of healthy donors and a C9ORF72-ALS patient at 4 weeks of adherent differentiation. SMN- and Gemin 2-double positive granules were counted as GEMs. Arrows indicate GEMs. Scale bar = 10 μm.

**Figure 5.**
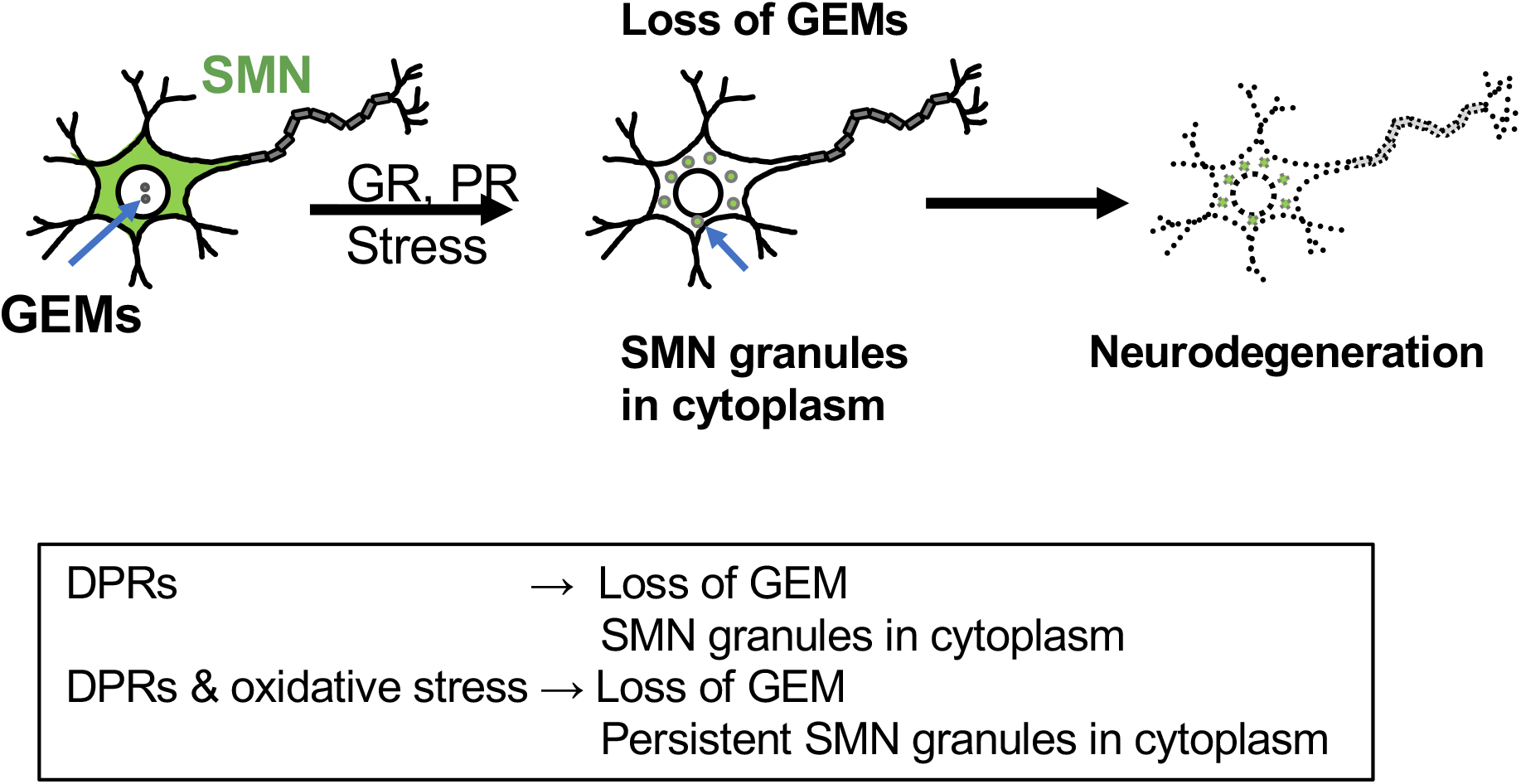
Graphical representation of the overall findings of this study. In normal cells, SMN protein forms phase-separated foci called GEMs in the nucleus and is diffusely distributed in the cytoplasm. Accumulation of *C9ORF72* poly(GR) and poly(PR) disturbs GEM formation and induces cytoplasmic granules harboring SMN and RNA-binding proteins. Poly(PR) expression combined with oxidative stress induces persistent cytoplasmic stress granules harboring SMN and RNA-binding proteins.

We have also previously shown that snRNPs extensively accumulate in MNs of postmortem tissues of ALS patients ^22^. To investigate whether snRNP accumulation occurred in MNs differentiated from the C9-ALS patient, we immunostained MNs with anti-Sm and anti-U1 70K antibodies. Anti-Sm antibody Y12 recognizes the methylated Sm core proteins of most snRNPs. U1 70K is a U1 snRNP-specific protein. There was no significant accumulation nor changes in their distribution (Supplementary Fig. S2), indicating that snRNP accumulation observed in the postmortem MNs of ALS patients seems to be the result of neurodegeneration. Together, these findings provide evidence for disruptions of GEMs in ALS patient-derived MNs harboring the *C9ORF72* mutation, suggesting low levels of SMN in the MNs of ALS patients and the actual involvement of GEM pathology in the degeneration of MNs in C9-ALS patients.

## Discussion

In this study, we showed that the expression of the *C9ORF72* DPR poly(GR) and, to a lesser extent, poly(PR) in HeLa cells results in the loss of GEM bodies. The analysis of MNs differentiated from iPSCs from an ALS patient harboring a *C9ORF72* mutation showed a decreased number of GEMs in MNs. Since the number of GEMs is an indicator of the abundance of functional SMN ^19^, these results indicate low levels of SMN in MNs of ALS patients harboring *C9ORF72* mutations. Furthermore, poly(GR) and poly(PR) induced the abnormal accumulation of SMN in the cytoplasm, especially after oxidative stress. Since SMN has a crucial role in snRNP assembly, low levels of SMN and abnormal cytoplasmic accumulation of SMN could lead to disturbed snRNP biogenesis. Taken together, we provide evidence that the level of functional SMN is low in MNs harboring the *C9ORF72* mutation.

What is the molecular mechanism to explain the loss of GEMs induced by poly(GR) or poly(PR)? Poly(GR) and poly(PR) have both been shown to interact with RNA-binding proteins and proteins with low complexity domains (also known as prion-like domains), thus damaging the assembly, dynamics, and functions of membrane-less phase-separated organelles ^31,32^. Since GEMs are a type of membrane-less organelle in the nucleus, it is likely that poly(GR) and poly(PR) directly alter the biophysical properties of GEMs by inhibiting liquid-liquid phase separation. GEMs often associate or overlap with Cajal bodies, where snRNAs undergo maturation, and some studies have shown that poly(GR) and poly(PR) impact the formation of Cajal bodies ^31,32^. The sequestration of SMN in cytoplasmic PABP granules may reduce the amount of SMN in the nucleus, resulting in an acceleration of a loss of GEMs. Alternatively, DPRs cause defects in nucleocytoplasmic transport ^36–39^, thus potentially leading to the retention of SMN in the cytoplasm. Other possible mechanisms leading to a low level of SMN are heterochromatin anomalies, dsRNA accumulation, and nucleolar stress induced by DPRs ^40,41^. These stresses might induce the degradation of SMN.

In this study, the sequestration of SMN in cytoplasmic PABP granules was observed in cells with poly(GR) or poly(PR). Poly(GR), poly(PR), and SMN are all able to bind to RNA^7^, so perhaps they interact in an RNA-dependent manner. Alternatively, several proteins that interact with poly(GR) or poly(PR) have been reported ^31,42–44^, and some of these might mediate the interaction with SMN and DPRs. Poly(GR) can bind ribosomal subunits and induce translational arrest ^40^, and SMN has also been found to associate with the ribosome and have a role in translation ^45,46^, indicating a possible link between DPRs and SMN.

*C9ORF72* mutations can cause disease by several different mechanisms. They can induce a loss of the C9ORF72 protein, a toxic gain of function from *C9ORF72* RNA accumulation or a toxic gain of function from the DPRs ^10^. Among these, poly(GR) and poly(PR) seem to be the most toxic to neurons ^7–9^. The pattern of poly(GR) accumulation seems to be the most correlated with neurodegeneration ^11,12^. Here, we showed that poly(GR) has stronger effects on GEM formation than poly(PR), that may explain the neurotoxicity of poly(GR) in ALS patients.

Here, we observed loss of GEMs but not abnormal snRNP accumulation in MNs differentiated from iPSCs of a C9ORF72-ALS patient at 4 weeks of differentiation. These observations may suggest that the loss of GEMs occurs at a relatively early stage and causes neurodegeneration and that snRNP accumulation is the result of the loss of SMN following neurodegeneration. The association of DPRs and U1 or U2 snRNP components ^31,42,44^ might contribute to the abnormal snRNPs observed in ALS patients.

In this report, we provided evidence that the level of functional SMN is low in MNs harboring a *C9ORF72* mutation. Importantly, increasing the level of SMN improves the survival of many types of human MNs, including MNs differentiated from iPSCs of SMA patients and ALS patients harboring *TDP-43* mutations, even with mutations in the ALS-related *SOD1* gene ^47^. These findings indicate the beneficial effect of increasing functional SMN in MNs and now suggest that this type of approach could be explored for *C9ORF72* mutations.

## Methods

### Cell culture and transfection

HeLa cells were maintained in DMEM/Ham's F-12 with L-glutamine medium (Thermo Fisher Scientific, USA) supplemented with 10% foetal bovine serum and a 0.5% penicillin-streptomycin mixed solution. In this study, a HeLa cell clone with a relatively large nucleus (HeLa-C3) was expanded and used. For the overexpression of DPRs, cells were transfected with plasmids using polyethylenimine "Max" (Polysciences, USA) with a standard method.

### iPSC culture and differentiation

iPSCs were maintained on mitomycin-C-treated SNL murine fibroblast feeder cells in 0.1% gelatine-coated tissue culture dishes in hESC medium. iPSCs were differentiated into spinal MNs as previously described ^33–35^. Briefly, iPSC colonies were detached using a dissociation solution (0.25% trypsin, 100 μg/ml collagenase IV (Thermo Fisher Scientific, USA), 1 mM CaCl_2_, and 20% KnockOut^TM^ Serum Replacement (KSR, Thermo Fisher Scientific, USA)) and cultured in suspension in bacteriological dishes in standard hESC medium. After the removal from SNL feeder cells, the cells were incubated for 1–2 hours in gelatine-coated dishes. On day 1, the medium was changed to human embryoid body (hEB) medium containing DMEM/F-12, 5% KSR, 2 mM L-glutamine, 1% MEM Non-Essential Amino Acids (NEAA), and 0.1 mM 2-mercaptoethanol with 300 nM LDN-193189 (Sigma-Aldrich, USA), 3 μM SB431542 (Tocris, UK), and 3 μM CHIR99021 (Focus Biomolecules, USA). On day 2, the medium was changed to fresh hEB medium containing 300 nM LDN-193189, 3 μM SB431542, 3 μM CHIR99021, and 1 μM retinoic acid (RA) (Sigma-Aldrich, USA). From day 4 to day 14, hEBs were cultured in hEB medium containing 1 μM RA and 1 μM purmorphamine (Calbiochem, Germany), and the medium was changed every 2–3 days. On day 14, hEBs were enzymatically dissociated into single cells using TrypLE Select (Thermo Fisher Scientific, USA). The dissociated cells were plated on growth factor-reduced Matrigel (33 × dilution, thin coating; Corning, USA)-coated dishes at a density of 2 × 10^4^ cells/well (96-well imaging plate, Greiner Bio-One, Austria) and cultured in MN medium (MNM) consisting of KBM Neural Stem Cell medium (Kohjin Bio, Japan) supplemented with 2% B27 supplement (Thermo Fisher Scientific, USA), 1% NEAA, 50 nM RA, 500 nM purmorphamine, 10 μM cyclic AMP (cAMP) (Sigma-Aldrich, USA), 10 ng/mL recombinant BDNF (R&D Systems, USA), 10 ng/mL recombinant GDNF (R&D Systems, USA), 10 ng/mL recombinant human IGF-1 (R&D Systems, USA), and 200 ng/mL L-ascorbic acid (Sigma-Aldrich, USA) for up to 4 weeks. Half of the medium was changed every 2–3 days. To visualize HB9-positive MNs, cells were infected with *HB9^e438^::Venus* lentivirus on day 4 as previously described ^33^. All the experimental procedures for the production and the use of iPSCs were approved by the ethics committee of the Aichi Medical University School of Medicine (approval number 14–004 and 2020-213).

### Immunocytochemical staining

Cells were fixed in 4% paraformaldehyde for 15–25 min at room temperature and permeabilized with 0.1% Triton-X. Nonspecific binding was blocked by incubation with 1% normal goat serum with 0.05% Triton X-100 for HeLa cells or 10% normal goat serum with 0.3% Triton X-100 for iPSC-derived MNs. Cells were then incubated with primary antibodies overnight at 4°C in PBS containing 1% BSA and 0.05% Triton X-100, rinsed with PBS three times, and incubated with Alexa Fluor-conjugated secondary antibodies (Thermo Fisher Scientific, USA) for 2 hours at room temperature. Nuclei were stained with 10 μg/ml Hoechst 33258 (Sigma-Aldrich, USA). The cells were then rinsed with PBS three times. The following antibodies were used: mouse anti-FLAG (F3165, Sigma, 1:200), rabbit anti-FLAG (F7425, Sigma, 1:200), rat anti-FLAG (NBP1-06712, Novus, 1:50), rabbit anti-PABP (ab21060, Abcam, 1:1000), mouse anti-PABP (P6246, Sigma, 1:500), mouse anti-SMN (IgG1, 610646, BD Biosciences, 1:250), mouse anti-Gemin2 (IgG2b, sc-32806, Santa Cruz, 1:50), goat anti-GFP (600-101-215, ROCKLAND, 1:500), rabbit anti-GFP (598, MBL, 1:500), mouse anti-HB9 (IgG1, 81.5C10, Developmental Studies Hybridoma Bank, 1:1000), mouse anti-Isl-1 (IgG2b, 39.4D5, Developmental Studies Hybridoma Bank, 1:400), and mouse anti-βIII-tubulin (IgG2a, MMS-435P, Covance, 1:2000). Alexa Fluor 488-, Alexa Fluor 555-, Alexa Fluor 594-, or Alexa Fluor 647-conjugated anti-mouse, anti-rabbit, anti-mouse IgG1, anti-mouse IgG2a, and anti-mouse IgG2b antibodies were used. HeLa cells were mounted using Dako Fluorescence Mounting Medium (S3023, Agilent Technology, USA).

Images of MNs differentiated from iPSCs and HeLa cells were obtained using a confocal microscope LSM700 (Carl Zeiss Microscopy, Germany) and a Keyence BZ-X700 microscope (Keyence, Japan), respectively. Images were analysed using Adobe Photoshop. For the counting of GEM in MNs differentiated from iPSCs, all Z-stack images at 2 μm intervals were analysed. SMN-positive granules in nuclei were counted as GEM in HeLa cells. SMN- and Gemin2-positive granules in nuclei were counted as GEM in MNs differentiated from iPSCs.

### RNA staining

Cells were prewashed with PBS and incubated with 500 mM SYTO RNASelect Green Fluorescent Cell Stain (S32703, Thermo Fisher Scientific, USA) for 20 min at 37°C, according to the instructions. Cells were then rinsed in medium and fixed with methanol for 10 min at −20°C, followed by immunostaining.

### Statistical analysis

Statistical analysis was performed using Prism 9 (GraphPad software, USA). Statistical tests included one-way and two-way ANOVA with Tukey’s multiple comparisons tests. All experiments are represented as the average ± standard error of the mean.

## Acknowledgements

This work was supported by Japan Society for the Promotion of Science KAKENHI Grant Numbers JP17K08280 and JP20K07016; the ALS Foundation, Japan ALS Association; the Ichihara International Scholarship Foundation; Grant-in-Aid for Research in Nagoya City University Grant Numbers 1 and 2022004 to HT. This work was also supported by grants from the Practical Research Project for Intractable Diseases and Research Center Network for Realization of Regenerative Medicine of the Japan Agency for Medical Research and Development (AMED) (JP19ek0109243 and JP20bm0804020) to YO.

## Author contributions

M.Y., I.N., and H.T. investigated and prepared figures 1-3. Y.K., I.N., K.O., Y.O., and H.T. investigated and prepared figures 4. Y.O. and H.T. wrote the main manuscript text. H.O., H.I., M.H., and Y.O. provided resources. All authors reviewed the manuscript.

