## Supplementary information for "*C9ORF72* dipeptide repeat proteins disrupt formation of GEM bodies and induce aberrant accumulation of survival of motor neuron protein"

**CONTENTS:**

**Supplementary Figure S1. Oxidative stress induces SMN accumulation in stress granules but not the loss of GEMs**

**Supplementary Figure S2. Distribution of snRNPs labelled with anti-Sm antibody and U1 70K protein in motor neurons derived from iPSC**

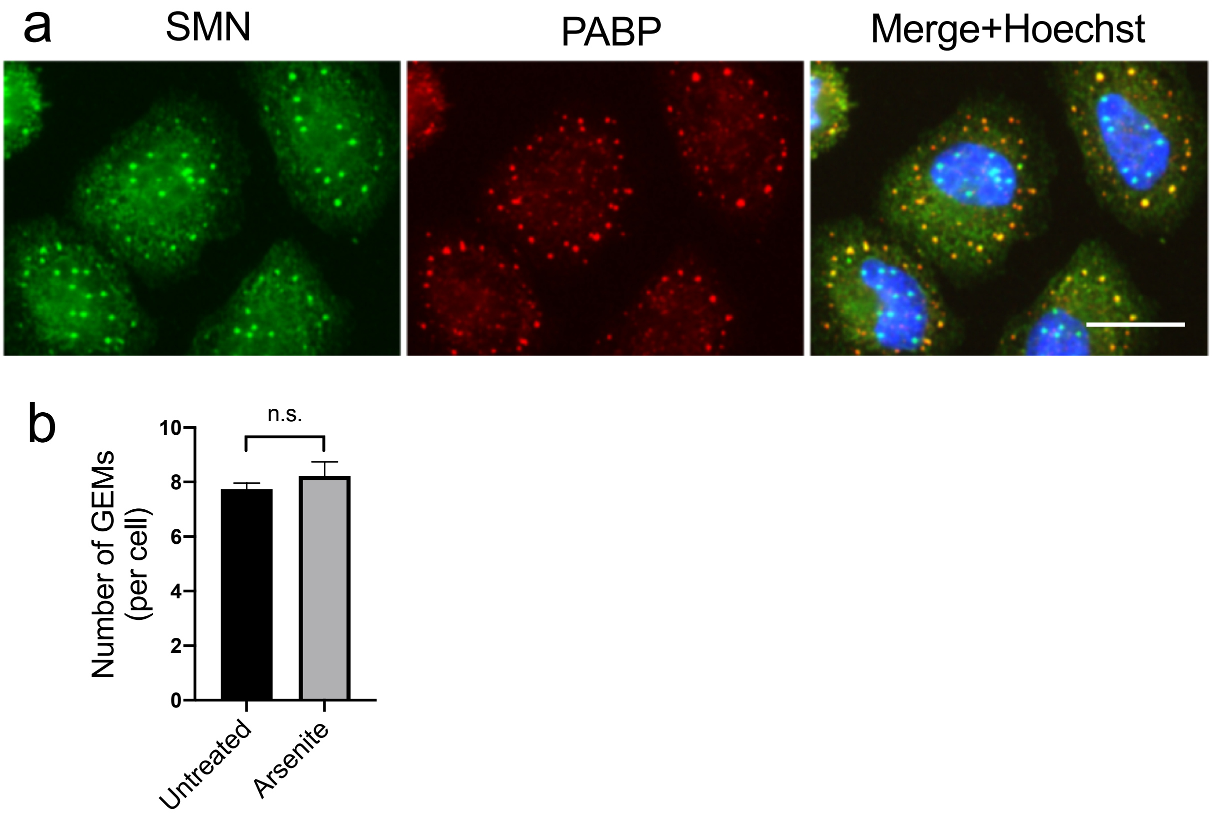


**Supplementary Figure S1. Oxidative stress induces SMN accumulation in stress granules but not the loss of GEMs**

(a) HeLa cells were treated with 500 μM sodium arsenite for 30 min to induce cytoplasmic stress granules, fixed, and immunostained with anti-PABP and anti-SMN antibodies. Scale bar = 20 μm.

(b) Quantification of the number of GEMs visualized in (a). Nuclear granules labelled with SMN antibody were counted as GEMs. More than 100 cells were counted. Unpaired t-test.

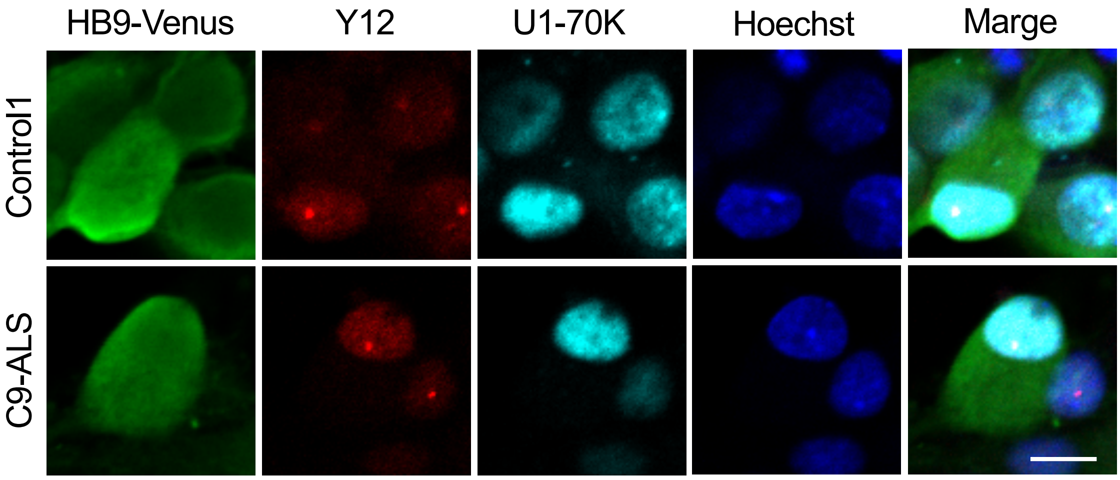


**Supplementary Figure S2. Distribution of snRNPs labelled with anti-Sm antibody and U1 70K protein in motor neurons derived from iPSCs**

Motor neurons differentiated from iPSCs of a healthy donor (Control1, upper row) or a C9ORF72-ALS patient (lower row) were stained with anti-GFP antibody, anti-Sm protein antibody (Y12), and anti-U1 70K antibody at 4 weeks of differentiation. Scale bar = 10 μm.
